# DrugPT: A Flexible Framework for Integrating Gene and Chemical Representations in Perturbation Modeling

**DOI:** 10.1101/2025.07.25.665130

**Authors:** Linchang Zhu, Yuanhanyu Luo

## Abstract

Accurately modeling the transcriptional response of cells to drug perturbations is critical for drug discovery and precision medicine. Here, we propose DrugPT, a novel Transformer-based framework for predicting gene expression changes upon drug treatment. DrugPT consists of three modular components: (1) a gene representation module that encodes pre-treatment expression profiles using GPT-derived embeddings; (2) a drug representation module that captures chemical structure information using a pre-trained GPT2 model; and (3) a Transformer-based prediction module that integrates gene and drug embeddings to predict perturbed gene expression profiles. Unlike previous approaches, DrugPT formulates the prediction task as a sequence modeling problem, leveraging pre-trained language models for both gene and drug modalities. We demonstrate the architecture of DrugPT and highlight its generalizable design for encoding heterogeneous biological and chemical information. This work introduces a unified modeling framework, laying the foundation for future developments in multimodal perturbation prediction tasks.

## Introduction

Modeling how drugs perturb gene expression in human cells is a fundamental challenge in systems biology, with broad implications for target identification, compound screening, and precision therapeutics^1^. Advances in high-throughput perturbation assays have enabled systematic profiling of transcriptional responses to thousands of chemical compounds. However, the ability to predict these responses from first principles given only a drug’s molecular structure and a cellular context remains limited^2^.

Recent studies have leveraged deep learning to predict perturbation outcomes, often by encoding drug structures using graph-based neural networks and modeling gene expression with fully connected layers or attention mechanisms^3-6^. While effective to some extent, these approaches often lack flexibility in representing complex multimodal relationships and fail to generalize across unseen drugs or cell types^7-10^.

The rapid expansion of high-throughput sequencing technologies has produced an enormous volume of perturbational data, yet the experimental exploration of the full chemical and cellular space remains intractable^10^. This limitation has spurred the development of computational methods, particularly deep learning models, to predict transcriptional responses and facilitate applications like drug repurposing^11,12^. Earlier models such as DeepCE^13^, which incorporates graph-based drug encoders, and DLEPS^14^, which uses deep neural networks to learn drug-induced expression shifts, have demonstrated the feasibility of such predictions. Generative approaches including PRnet and TranSiGen^15,16^ have also shown success in reconstructing perturbation signatures using variational autoencoder frameworks^17^.

Here, we present DrugPT, a new Transformer-based framework for drug perturbation modeling. Inspired by advances in language modeling, DrugPT employs pretrained GPT-style encoders to generate contextual embeddings for both gene expression profiles and drug structures. By framing the gene-drug interaction as a sequence modeling task, our model captures higher-order dependencies and compositional semantics between cellular states and chemical perturbations.

DrugPT introduces a modular and extensible design with three key components: (1) a gene representation module that transforms pre-treatment expression into a contextualized embedding space; (2) a drug representation module built upon GPT2-derived embeddings of chemical structures; and (3) a Transformer module that integrates these embeddings to predict post-treatment expression profiles.

Rather than focusing on architecture-level innovations, this study emphasizes the formulation of a unified, and generalizable modeling framework for drug perturbation prediction. DrugPT establishes a foundation for future integration of multi-omic, temporal, and spatial modalities into next-generation perturbation models.

## Results

### A modular Transformer framework for perturbation response prediction

To address the challenge of modeling gene expression responses to drug perturbations, we developed DrugPT, a unified framework that integrates molecular and cellular information using a Transformer-based architecture. DrugPT comprises three key components designed to separately encode pre-treatment cell states and chemical structures before integrating them for response prediction (Figure 1).

**Fig. 1:**
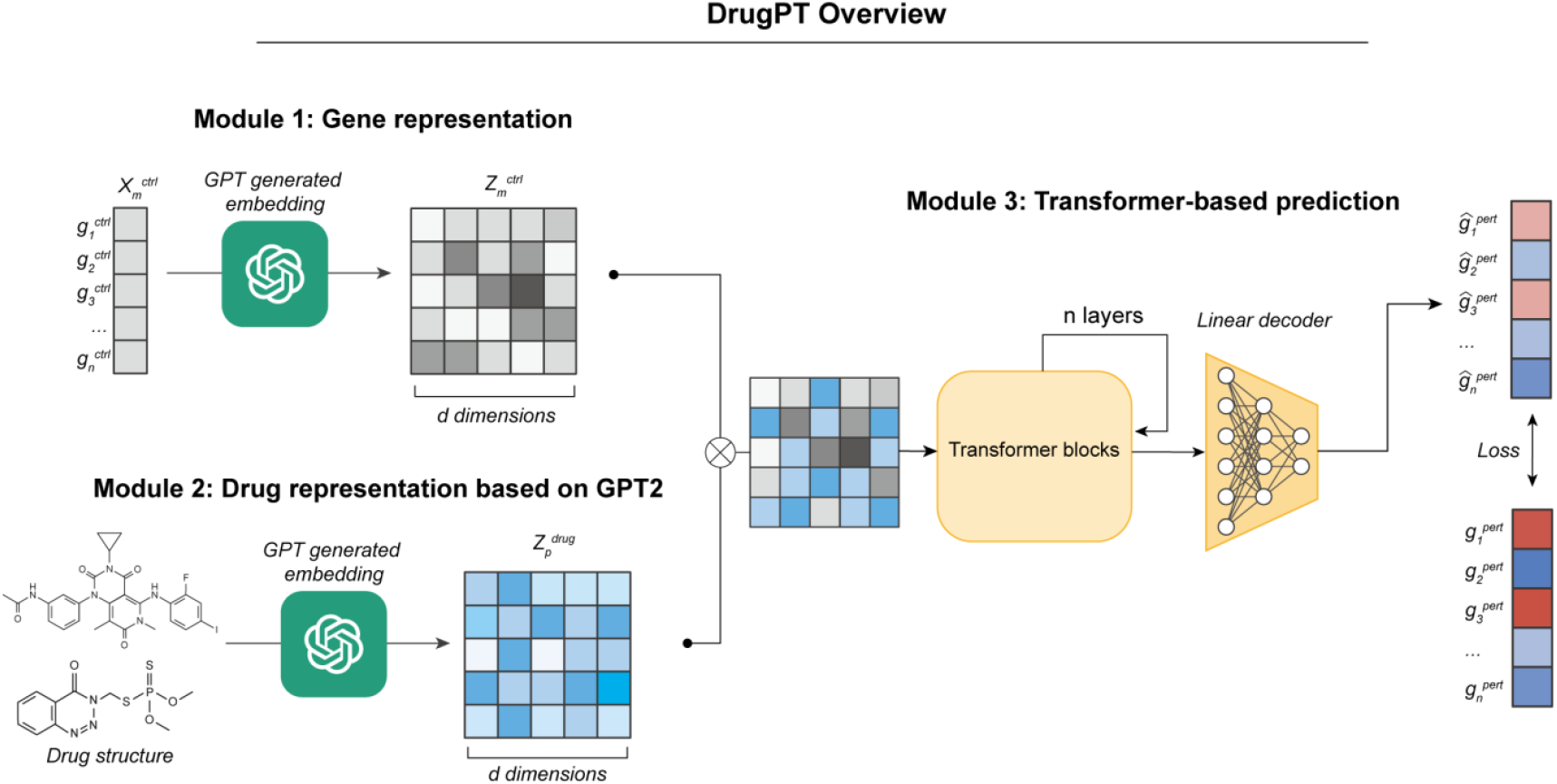
Overview of the DrugPT architecture. DrugPT is a modular Transformer-based model designed to predict drug-induced gene expression changes by integrating gene and drug representations. The framework comprises three main components. First, a gene representation module encodes the pre-treatment expression profile of each cell using a GPT-derived embedding model, producing a contextualized gene embedding matrix Z_ctrl from the original expression input X_ctrl. Second, a drug representation module transforms the chemical structure of a compound into a vectorized embedding Z_drug using a GPT2-based encoder trained on molecular graphs. The gene and drug embeddings are then fused and fed into a stack of Transformer blocks, which model the interaction between cellular context and chemical perturbation. Finally, a linear decoder predicts the perturbed expression profile g_hat_pert, and the model is trained to minimize the difference between predicted and observed gene expression responses.

The first component is a gene representation module, which transforms the pre-treatment gene expression profile of a cell into a contextualized embedding. Instead of relying on handcrafted gene features or shallow encoders, we utilize a pretrained GPT-style model to capture higher-order relationships among genes based on their co-expression structure. Each input profile is converted into a sequence of gene embeddings, preserving positional and distributional dependencies that may inform downstream responses.

The second component is a drug encoder, which employs a GPT2-based architecture trained on drug structure to model the structural properties of chemical compounds. This encoder generates an embedding that is dimensionally aligned with the gene representation, enabling seamless fusion. Unlike graph convolutional encoders that require explicit molecular graphs, our language-model-based drug encoder naturally captures substructure patterns and global molecular semantics.

The final component is a Transformer integration module that receives the concatenated gene and drug embeddings as input. This module learns to model the complex interplay between cellular context and chemical perturbation by attending across both modalities. A linear decoder then maps the integrated embedding back to the predicted post-treatment expression profile.

Together, these components form a flexible and extensible architecture that can be trained end-to-end. DrugPT reframes perturbation modeling as a multimodal sequence prediction problem, making it amenable to transfer learning, pretraining, and future integration of additional modalities such as genetic perturbations or time-series measurements. Unlike previous models that often entangle drug and gene representations in early stages, DrugPT explicitly separates and then aligns them within a controlled Transformer context, allowing more interpretable and modular design.

### DrugPT generalizes to unseen compounds in a drug-holdout evaluation

We first evaluated the ability of DrugPT to generalize to unseen chemical compounds using a drug-holdout prediction task. In this setting, the model was trained on 80% of the compounds and evaluated on the remaining 20% that were not observed during training (Figure 2a). This setup simulates the practical challenge of predicting responses to novel drugs based on prior transcriptomic and chemical data.

**Fig. 2:**
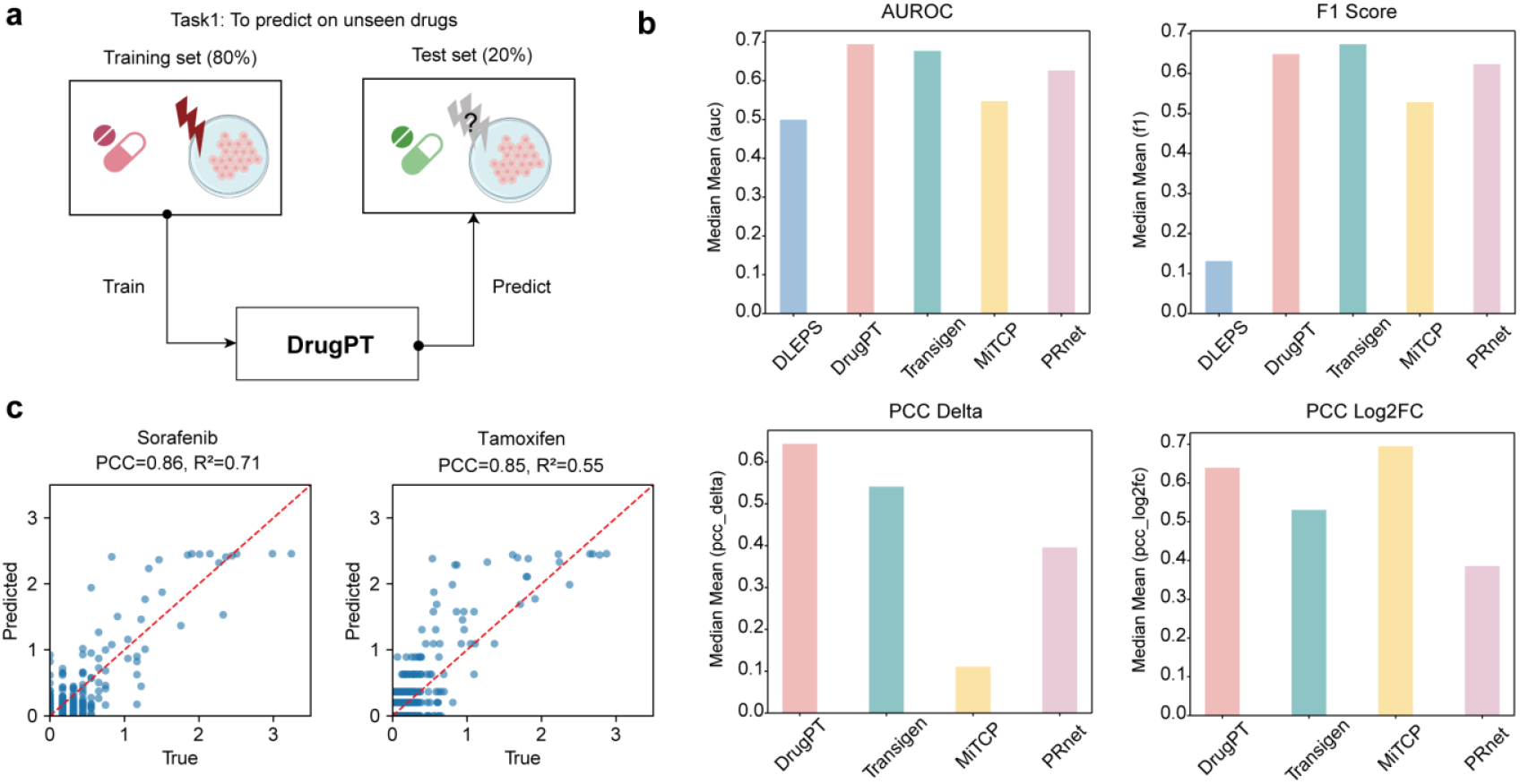
Evaluation of DrugPT on the drug-holdout prediction task. a, Schematic of the experimental setup for Task 1. The model is trained on gene expression profiles from 80% of the compounds and evaluated on the remaining 20% of drugs not seen during training. b, Comparison of DrugPT with four baseline models (DLEPS, Transigen, MitCP, and PRnet) using four evaluation metrics: AUROC, F1 score, PCC based on delta expression, and PCC based on log2 fold change. DrugPT achieves strong and consistent performance, particularly on delta-based Pearson correlation. c, Scatter plots showing the predicted versus true post-treatment gene expression levels for two representative held-out drugs, Sorafenib and Tamoxifen. DrugPT predictions show high concordance with experimental values, with Pearson correlation coefficients of 0.86 and 0.85, respectively.

To benchmark performance, we compared DrugPT with four representative models: DLEPS, Transigen, MitCP, and PRnet. DrugPT achieves competitive performance across a range of evaluation metrics. On classification tasks such as AUROC and F1 score, DrugPT performs comparably to or better than existing approaches (Figure 2b). On regression-based metrics that directly assess expression accuracy, DrugPT attains the highest Pearson correlation coefficient (PCC) when evaluated on delta expression profiles, suggesting its strong capacity to model expression change rather than absolute levels.

We further visualized predictions on two representative held-out drugs, Sorafenib and Tamoxifen (Figure 2c). In both cases, DrugPT predictions aligned well with observed post-treatment profiles, with PCC values of 0.86 and 0.85, respectively. Although the model does not outperform all baselines on every metric, these results highlight the framework’s ability to capture perturbational effects for unseen compounds in a realistic setting. We note that DrugPT’s architecture does not require drug-specific fine-tuning and operates in a zero-shot manner for unseen inputs, offering practical advantages for future applications.

### DrugPT maintains performance across unseen cellular contexts

To further assess the generalization capability of DrugPT, we conducted a cell line-holdout evaluation, where the model was trained on transcriptional responses from a subset of cell lines and evaluated on held-out lines not seen during training (Figure 3a). This setup mimics the common real-world challenge of predicting drug responses in new or poorly profiled cellular systems.

**Fig. 3:**
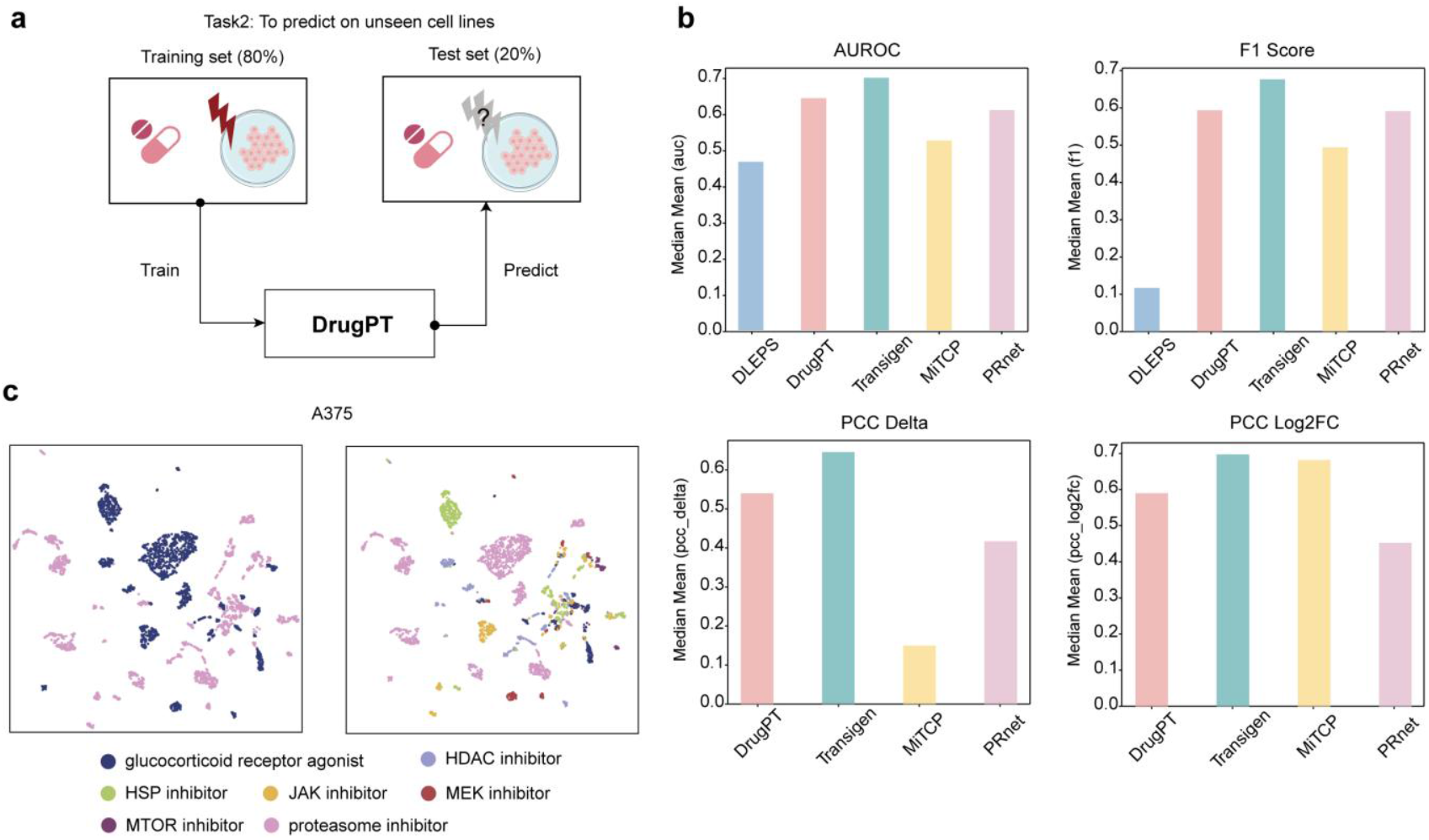
Evaluation of DrugPT on the cell line-holdout prediction task. a, Overview of Task 2 setup, where the model is trained on 80% of cell lines and tested on the remaining 20% that are not included during training. This setting assesses the model’s ability to generalize across cellular contexts. b, Benchmark comparison across multiple models using AUROC, F1 score, Pearson correlation on delta expression, and Pearson correlation on log2 fold change. DrugPT maintains stable performance on both classification and regression tasks when evaluated on unseen cell lines. c, UMAP visualization of true (left) and predicted (right) expression profiles for compound perturbations in a representative test cell line. Drug classes are colored by annotated mechanism of action, demonstrating the model’s ability to recover biologically meaningful structure in the predicted responses.

We benchmarked DrugPT against several representative models using the same evaluation metrics as in the drug-holdout task. As shown in Figure 3b, DrugPT achieves consistently strong performance across both classification metrics (AUROC and F1 score) and regression metrics (Pearson correlation based on delta expression and log2 fold change). Although some competing methods attain similar or slightly higher scores on specific metrics, DrugPT demonstrates robust and balanced performance without task-specific tuning.

To visualize the quality of predicted expression responses, we projected both ground truth and model-predicted profiles for a held-out cell line using UMAP (Figure 3c). The predicted signatures preserve the clustering of compounds with similar mechanisms of action, suggesting that DrugPT not only approximates global expression changes but also retains biologically relevant structure. These results indicate that DrugPT is capable of generalizing beyond training cell lines and may support applications in novel or rare cellular contexts..

## Discussion

In this study, we introduced DrugPT, a modular Transformer-based framework for predicting gene expression responses to chemical perturbations. By combining GPT-derived embeddings of both gene expression and molecular structure, DrugPT formulates the perturbation prediction task as a multimodal sequence modeling problem. This design allows the model to integrate heterogeneous biological and chemical information in a flexible and extensible manner.

Through systematic evaluations on both drug-holdout and cell line-holdout settings, we show that DrugPT achieves competitive performance across multiple metrics and tasks. The model demonstrates a stable ability to generalize to previously unseen compounds and cellular contexts without the need for retraining or task-specific tuning. Moreover, qualitative analyses suggest that DrugPT can capture biologically meaningful structures in the predicted transcriptional responses, including the grouping of compounds by mechanism of action.

Despite these encouraging results, we acknowledge several limitations. First, our current implementation relies on pretrained language models for molecular and expression representation, which may not fully capture the mechanistic underpinnings of gene regulatory networks or drug-target interactions. Second, the model operates at the bulk transcriptome level and does not account for cell type heterogeneity or temporal dynamics, which are increasingly important in perturbation studies. Finally, while DrugPT provides a generalizable framework, further investigation is needed to assess its utility in real-world applications such as drug repurposing, combinatorial therapy design, or cross-species translation.

Looking forward, several extensions are possible. The modular structure of DrugPT allows for the incorporation of additional input modalities, such as genetic perturbations, pathway annotations, or single-cell transcriptomic data. Future work may also explore more fine-grained training objectives, such as contrastive learning or causal modeling, to improve biological interpretability. Ultimately, we hope this framework can serve as a foundation for building more versatile and biologically grounded perturbation models.

## Author contributions

Y. Luo and L. Zhu designed the framework and performed data. Y. Luo and L. Zhu wrote the manuscript with input from all authors. All authors read and approved the final manuscript.

## Competing interests

The authors declare that they have no competing interests.

**Figure Legends**

## Methods

### Data sources and preprocessing

We utilized publicly available gene expression data from the LINCS L1000 dataset, which provides transcriptomic profiles of human cell lines under various small molecule perturbations. Each sample contains a vector of expression values for 978 landmark genes, measured before and after drug treatment. For each compound cell pair, we extracted the pre-treatment expression profile as model input, and the post-treatment profile as the target to be predicted. Drug chemical structures were processed using a pretrained GPT2 tokenizer. Cell lines and drugs were independently split into training and testing subsets to create two evaluation settings.

### Gene representation module

To encode pre-treatment gene expression profiles, we employed a GPT-derived embedding strategy. Each input vector, representing the expression levels of landmark genes in a specific cell state, was mapped into a contextualized embedding space using a pretrained sequence encoder. This approach captures co-expression dependencies among genes and provides a dense, learned representation for downstream integration. The resulting gene embedding matrix maintains the dimensionality alignment required for fusion with drug representations.

### Drug representation module

Chemical compounds were embedded using a pretrained GPT2-based encoder. The tokenizer converts the chemical compounds into token sequences, which are then passed through transformer layers to obtain fixed-length embeddings. The output embedding captures both local substructure features and global molecular semantics, allowing for the modeling of structure activity relationships without explicit molecular graphs. Drug embeddings were reshaped to align with the dimension of the gene embedding matrix, enabling direct integration in the subsequent Transformer module.

### Multimodal integration and prediction

The gene and drug embeddings were concatenated and input into a series of Transformer blocks. Each block contains multi-head self-attention and feed-forward sublayers, with layer normalization and residual connections applied throughout. The Transformer integrates information across the combined sequence, allowing the model to learn gene drug interactions in a context-aware manner. The final hidden states were passed through a linear decoder to predict the perturbed gene expression vector, corresponding to the post-treatment profile.

### Model training

The model was trained to minimize the mean squared error (MSE) between the predicted and ground-truth post-treatment expression vectors. Optimization was performed using the Adam optimizer with a learning rate of 1e-4 and a batch size of 64. A dropout rate of 0.1 was applied to improve generalization. The model was trained for up to 100 epochs with early stopping based on validation loss. All training was conducted using PyTorch on NVIDIA GPUs.

### Baseline models and evaluation metrics

To benchmark the performance of DrugPT, we compared it against several existing perturbation modeling approaches, including DLEPS, Transigen, PRnet, and MitCP. For fair comparison, we used publicly available implementations and followed the original training protocols for each method. Model performance was assessed using both classification and regression metrics. For classification, we computed AUROC and F1 score based on binarized gene regulation states. For regression, we calculated the Pearson correlation coefficient between predicted and true expression values, both for delta expression and log2 fold change. Additionally, we used UMAP to visualize predicted expression profiles and assess the preservation of biological structure in compound clustering.

### Implementation details

All models were implemented in Python using PyTorch. The GPT2 tokenizer and encoder were based on the HuggingFace Transformers library. Code for data processing, model training, and evaluation will be made available upon publication to ensure reproducibility.

## Notes

### Competing Interest Statement

The authors have declared no competing interest.

## References

1. Zhang K, Yang X, Wang Y, et al. Artificial intelligence in drug development. Nat Med. Jan 2025;31(1):45–59. doi:10.1038/s41591-024-03434-4

2. Moffat JG, Rudolph J, Bailey D. Phenotypic screening in cancer drug discovery - past, present and future. Nat Rev Drug Discov. Aug 2014;13(8):588–602. doi:10.1038/nrd4366

3. Pink R, Hudson A, Mouries MA, Bendig M. Opportunities and challenges in antiparasitic drug discovery. Nat Rev Drug Discov. Sep 2005;4(9):727–40. doi:10.1038/nrd1824

4. Siqueira-Neto JL, Wicht KJ, Chibale K, Burrows JN, Fidock DA, Winzeler EA. Antimalarial drug discovery: progress and approaches. Nat Rev Drug Discov. Oct 2023;22(10):807–826. doi:10.1038/s41573-023-00772-9

5. Renaud JP, Chung CW, Danielson UH, et al. Biophysics in drug discovery: impact, challenges and opportunities. Nat Rev Drug Discov. Oct 2016;15(10):679–98. doi:10.1038/nrd.2016.123

6. Moffat JG, Vincent F, Lee JA, Eder J, Prunotto M. Opportunities and challenges in phenotypic drug discovery: an industry perspective. Nat Rev Drug Discov. Aug 2017;16(8):531–543. doi:10.1038/nrd.2017.111

7. Lamb J, Crawford ED, Peck D, et al. The Connectivity Map: using gene-expression signatures to connect small molecules, genes, and disease. Science. Sep 29 2006;313(5795):1929–35. doi:10.1126/science.1132939

8. Subramanian A, Narayan R, Corsello SM, et al. A Next Generation Connectivity Map: L1000 Platform and the First 1,000,000 Profiles. Cell. Nov 30 2017;171(6):1437–1452 e17. doi:10.1016/j.cell.2017.10.049

9. Schenone M, Dancik V, Wagner BK, Clemons PA. Target identification and mechanism of action in chemical biology and drug discovery. Nat Chem Biol. Apr 2013;9(4):232–40. doi:10.1038/nchembio.1199

10. Sirota M, Dudley JT, Kim J, et al. Discovery and preclinical validation of drug indications using compendia of public gene expression data. Sci Transl Med. Aug 17 2011;3(96):96ra77. doi:10.1126/scitranslmed.3001318

11. Douglass EF, Jr., Allaway RJ, Szalai B, et al. A community challenge for a pancancer drug mechanism of action inference from perturbational profile data. Cell Rep Med. Jan 18 2022;3(1):100492. doi:10.1016/j.xcrm.2021.100492

12. Singh J, Petter RC, Baillie TA, Whitty A. The resurgence of covalent drugs. Nat Rev Drug Discov. Apr 2011;10(4):307–17. doi:10.1038/nrd3410

13. Pham TH, Qiu Y, Zeng J, Xie L, Zhang P. A deep learning framework for high-throughput mechanism-driven phenotype compound screening and its application to COVID-19 drug repurposing. Nat Mach Intell. Mar 2021;3(3):247–257. doi:10.1038/s42256-020-00285-9

14. Tong X, Qu N, Kong X, et al. Deep representation learning of chemical-induced transcriptional profile for phenotype-based drug discovery. Nat Commun. Jun 25 2024;15(1):5378. doi:10.1038/s41467-024-49620-3

15. Zhu J, Wang J, Wang X, et al. Prediction of drug efficacy from transcriptional profiles with deep learning. Nat Biotechnol. Nov 2021;39(11):1444–1452. doi:10.1038/s41587-021-00946-z

16. Qi X, Zhao L, Tian C, et al. Predicting transcriptional responses to novel chemical perturbations using deep generative model for drug discovery. Nat Commun. Oct 26 2024;15(1):9256. doi:10.1038/s41467-024-53457-1

17. Rees MG, Seashore-Ludlow B, Cheah JH, et al. Correlating chemical sensitivity and basal gene expression reveals mechanism of action. Nat Chem Biol. Feb 2016;12(2):109–1

